# OOPS: Object-Oriented Polarization Software for analysis of fluorescence polarization microscopy images

**DOI:** 10.1101/2023.12.05.570241

**Authors:** William F. Dean, Tomasz J. Nawara, Rose M. Albert, Alexa L. Mattheyses

## Abstract

Most essential cellular functions are performed by proteins assembled into larger complexes. Fluorescence Polarization Microscopy (FPM) is a powerful technique that goes beyond traditional imaging methods by allowing researchers to measure not only the localization of proteins within cells, but also their orientation or alignment within complexes or cellular structures. FPM can be easily integrated into standard widefield microscopes with the addition of a polarization modulator. However, the extensive image processing and analysis required to interpret the data have limited its widespread adoption. To overcome these challenges and enhance accessibility, we introduce OOPS (Object-Oriented Polarization Software), a MATLAB-based analysis tool tailored for FPM data. This work highlights the distinctive features of our software, which empower researchers to efficiently manage large datasets; detect and analyze individual structures; conduct population assessments based on morphology, intensity, and polarization-specific parameters; and create publication-quality visualizations, all within a user-friendly graphical interface. Importantly, OOPS is adaptable to various sample types, labeling techniques, and imaging setups, facilitating in-depth analysis of diverse polarization-sensitive samples. Here, we demonstrate the power and versatility of our approach by applying OOPS to FPM images of both punctate and filamentous structures. OOPS is freely available under the GNU GPL 3.0 license and can be downloaded at https://github.com/Mattheyses-Lab/OOPS.

## Introduction

Fluorescence polarization microscopy (FPM) encompasses a powerful set of techniques that enable measurements of macromolecular order and orientation in biological systems (1). FPM techniques come in many flavors (2), all of which are based upon the anisotropic nature of fluorescence absorption and emission (3). It is possible to study the organization of fluorescently labeled molecules microscopically by either modulating the orientation of the excitation polarization (4-7), separating the emission into polarized components (8, 9), or both (10, 11). We focus here on the excitation-resolved modality, which can be configured on a standard widefield microscope by simply placing a polarization modulating device in the illumination path.

Excitation-resolved FPM has provided novel insights into the architecture of cell-cell junctions (6, 7), the ordering of septin filaments in budding yeast (4, 5, 12), and the orientation of integrin traction forces (13), among others. However, full utilization of this technique has been hindered by the substantial image processing and mathematical analysis required to interpret the acquired data. As a result, its adoption has remained limited to a small number of specialized laboratories, often reliant on custom software solutions tailored to specific imaging setups or specimen geometries and frequently lacking comprehensive descriptions, thereby impeding their accessibility to the broader scientific community. Recognizing this need, multiple high-quality open-source software tools have emerged that are capable of processing excitation-resolved FPM data (14, 15). However, it remains challenging to manage large FPM datasets and perform object-based analyses.

Here, we present OOPS (Object-Oriented Polarization Software), an advanced object-oriented image analysis software tailored for excitation-resolved FPM. By combining efficient data management; flexible image segmentation; extensive object feature extraction; effortless object selection, filtering, and labeling; automated dipole orientation analysis; and highly customizable, publication-quality visualization, our software empowers researchers to unlock the full potential of excitation-resolved FPM. In the interest of accessibility, all of the processing, analyses, and visualizations are housed within a convenient graphical user interface. We anticipate that this software will facilitate wider adoption of FPM, enhance analysis capabilities, and streamline workflows, ultimately driving scientific discoveries.

## Design and implementation

OOPS is a GUI-based platform for object-based image analysis of fluorescence polarization microscopy images. The software is implemented in MATLAB and can be freely downloaded from GitHub. Usage requires an installation of MATLAB R2023b, as well as several MATLAB toolboxes, which are described in the documentation included with the software. OOPS uses BioFormats to import image data and can handle any open-source or proprietary microscopy image format supported by BioFormats (16).

### Expressions used to retrieve FPM statistics

Fluorescence Polarization Microscopy (FPM) aims to retrieve order and orientation statistics for a molecule or molecular complex. The core calculations are based on the anisotropic nature of fluorescence excitation with plane-polarized light. In a radiatively dominated regime, fluorescence intensity (*I*) is proportional to absorption probability such that:

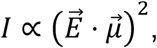

where 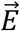 is a vector describing the amplitude and orientation of the excitation field and 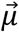 is the absorption transition moment of the fluorophore (17). For a plane-polarized excitation beam propagating along a fixed optical axis (z), 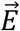 lies in the sample plane (x-y) and is described by its azimuthal angle (ω). The orientation of 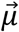 is described by its polar (*θ*) and azimuthal (*φ*) angles. Here, the azimuthal angle is the angle between the positive x-axis and the orthogonal projection of 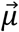 onto the x-y plane, while the polar angle is the angle between 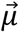 and the z-axis. The fluorescence intensity measured using a particular excitation polarization (*I*_*φ*_) is:

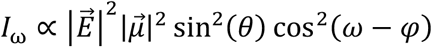

Thus, for a single, static fluorophore, *I*_*ω*_ varies sinusoidally with ω, and the orientation of 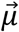 can be determined by making multiple intensity measurements while rotating 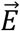 . For many fluorophores within a single diffraction-limited spot—assuming the rotational correlation time of the fluorophore is much slower than the fluorescence lifetime—the peak-to-peak amplitude of the sinusoid reflects the in-plane orientational order of the absorbing molecules, which we refer to as “order”. The phase of the sinusoid gives the average azimuthal direction of the fluorophore transition dipole moments, which we refer to interchangeably as “azimuth” or *α*. The order and azimuth can both be calculated using pixelwise image arithmetic from a series of images captured using excitation polarizations of 0°, 45°, 90°, and 135°:

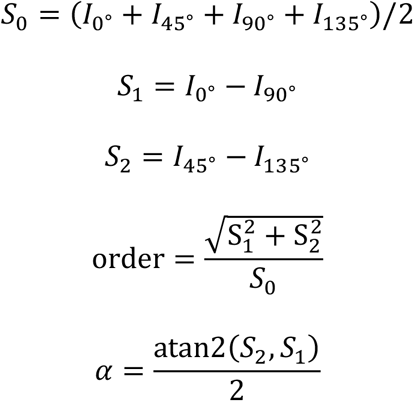

### Data requirements

Similar to many FPM implementations, our setup involves acquiring a stack of four fluorescence images using excitation polarizations of 0°, 45°, 90°, and 135° (FPM stack). However, one of the major hurdles to broad adoption of FPM is that researchers employ disparate sets of nuanced correction and calibration procedures tailored to their specific microscope setups, acquisition settings, and downstream analyses. As a result, there does not exist a standard set of pre-processing operations, even when the excitation polarizations used to acquire the data are identical. By contrast, once the raw data have been sufficiently processed, the equations used to retrieve the order and azimuth are extremely similar. For this reason, the only calibration procedure employed by OOPS is a standard flat-field correction, which is itself optional. If desired, users can upload one or more flat-field calibration stacks captured using the same excitation polarizations as the FPM stack. However, the only required input is a single FPM stack, which can be pre-processed in any manner.

While initially designed for excitation-resolved data, OOPS can easily be extended to emission-resolved setups, assuming the emission is split into 0°, 45°, 90°, and 135° components. In this case, or in the case of more nuanced acquisition setups, it may be desirable to change how the order statistic is calculated. To that end, we provide `CustomFPMStatistic.m`, a configuration class which the user can instantiate by providing: the name of the custom statistic, the display name to be used in GUI elements and plots, the possible range of values, and a handle to a function that accepts a single FPM stack and returns the pixelwise calculation of the custom statistic. An instance of the class is saved as a configuration file and read by OOPS during startup. The software will dynamically add the user-defined properties to the data classes and seamlessly integrate the new statistic into the interface. Importantly, in addition to defining custom order parameters, this functionality also provides a way to add custom correction and calibration procedures to the software.

### Data structure

To facilitate simultaneous analysis of datasets containing multiple experimental conditions or replicates, OOPS employs a hierarchical data structure (Fig 1A). After opening the software, users create a “project” organized into “groups”, each corresponding to a different set of experimental conditions. Groups are composed of one or more “images”—each storing the raw FPM stack and any associated output data for a given technical replicate—along with one or more optional flat-field normalization stacks. Once the data have been loaded, analysis results in the construction of “objects”, which contain all of the extracted features and output statistics for individual structures detected in an image (Fig 1B).

**Figure 1.**
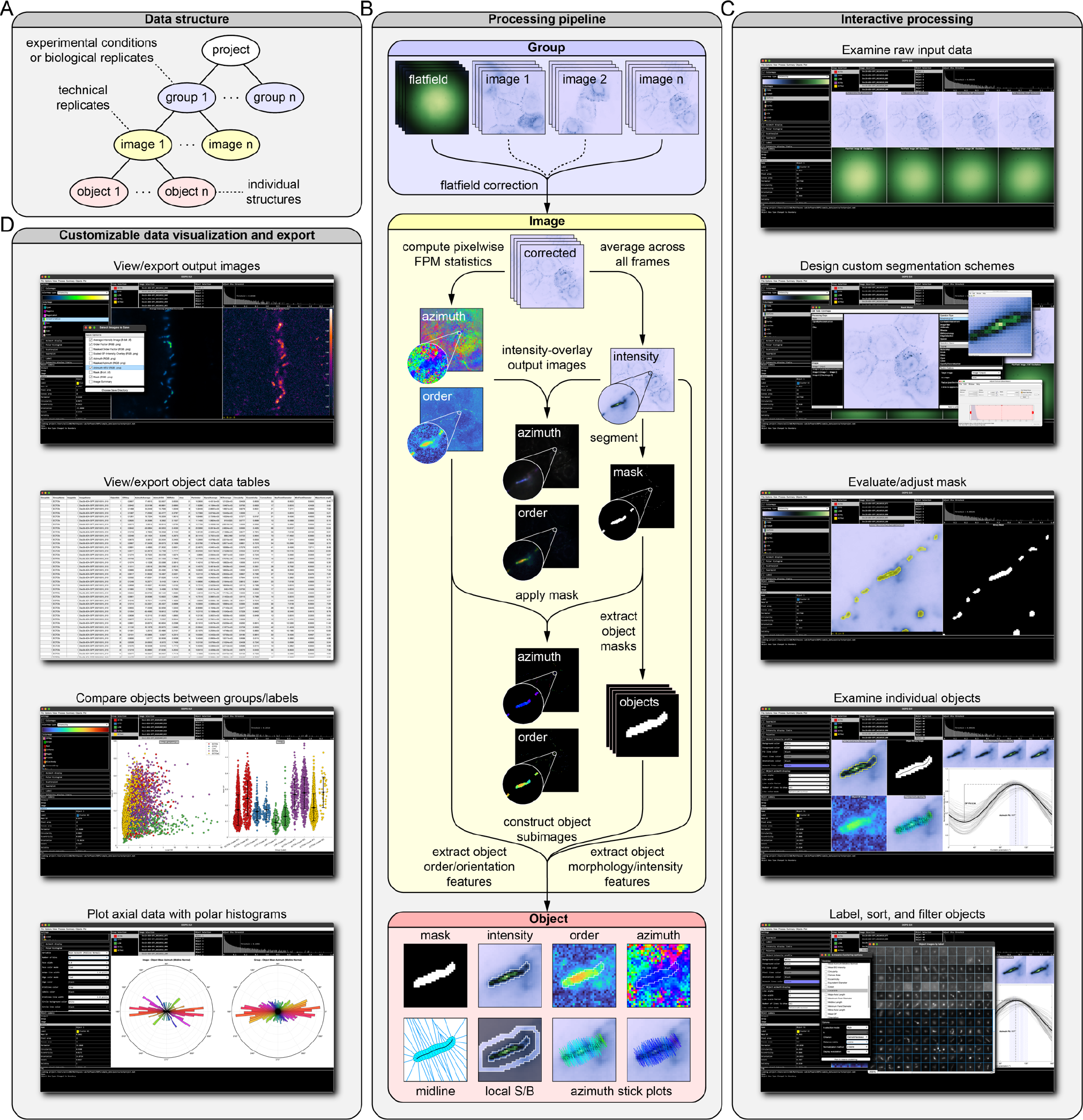
Overview of Object-Oriented Polarization Software (OOPS). (*A*) Hierarchical data structure used by the software. Projects are organized into “groups” representing biological replicates or experimental conditions, each containing a certain number of “images”. Each image contains “objects”, which store properties and statistics calculated for individual structures in the image. (*B*) Simplified processing pipeline for a single “image”, which includes flat-field correction, segmentation, calculation of FPM statistics, and object feature extraction. (*C*) Screenshots from the OOPS GUI showing examples of interactive data processing including examination of data, design of custom segmentation schemes, adjustment of image masks, and object manipulation. (*D*) Screenshots showing examples of the different image, plot, and data table visualizations available in the software.

### Interactive data analysis, visualization, and export

Users interactively guide the data through each processing and analysis step with a flexible level of supervision (Fig 1C). Using default settings, users can quickly perform flat-field corrections, calculate pixelwise order and orientation statistics, segment images into objects, and extract object features for multiple images with only a few clicks. Alternatively, users can create custom segmentation schemes; adjust image masks; select, view, and label objects; and perform property-based object filtering, sorting, and grouping for more detailed analyses. Once the analyses are complete, users will visualize data with a wide variety of customizable plots and images (Fig 1D), which can be exported or copied directly from the software. Detailed object data tables can be exported for use in other plotting or statistics software. At any point during the analysis, the project can be saved, closed, and then reopened at a later stage to continue the analysis.

## Results

OOPS was designed to democratize Fluorescence Polarization Microscopy (FPM) analysis for cell biology laboratories. The software is insensitive to both sample morphology and labeling strategy, a versatility we showcase here through the analysis of two distinct datasets: transmembrane adhesion receptors labeled with genetically encoded fluorophores and cytoskeletal filaments labeled with small molecule fluorophores. In this section, we detail how OOPS equips users with a diverse array of customization options to effectively visualize image data. Moreover, we illustrate the transformative potential of an object-based analysis approach, demonstrating its capacity to elevate data quality, unveil hidden relationships, and facilitate object clustering for more in-depth analyses. Lastly, we explore object-based azimuth orientation analysis as a tool to reveal underlying geometric features and quantify changes in orientation.

### Image types and visualization options

The software enables users to export image data with a wide variety of customization options. To illustrate different types of output images, we fluorescently labeled the F-actin cytoskeleton of COS-7 cells with Alexa Fluor 488 (AF488) phalloidin—an ideal sample for FPM due to the fact that light absorption is strongly polarized along the direction parallel to the filaments (18). The main FPM-specific image outputs are the pixelwise order and azimuth maps. These, along with the average intensity image, can be directly exported for downstream analyses. In addition to the raw, grayscale output, the software enables the export of special output types that combine multiple statistics into a single image, each of which offer unique benefits.

For example, the order-intensity overlay is constructed by merging the order and intensity images, with the intensity image acting as opacity mask (Fig 2A). This method is effective at dampening the noisy background regions of the order image, while not obscuring the sample itself. The contrast of the intensity and order channels can be set independently by the user, and the image can be colorized using a large selection of lookup tables (LUTs). The azimuth data are inherently more difficult to display in image form, due to the angular nature of the measurements. By default, the azimuth data are wrapped to the range [0°,180°] and displayed with a circular LUT where values of 0° and 180° map to the first and last colors in the LUT, respectively (Fig 2B). Similar to the order image, the azimuth image can be displayed using the intensity image as an opacity mask (not shown), which offers the same benefits mentioned above.

**Figure 2.**
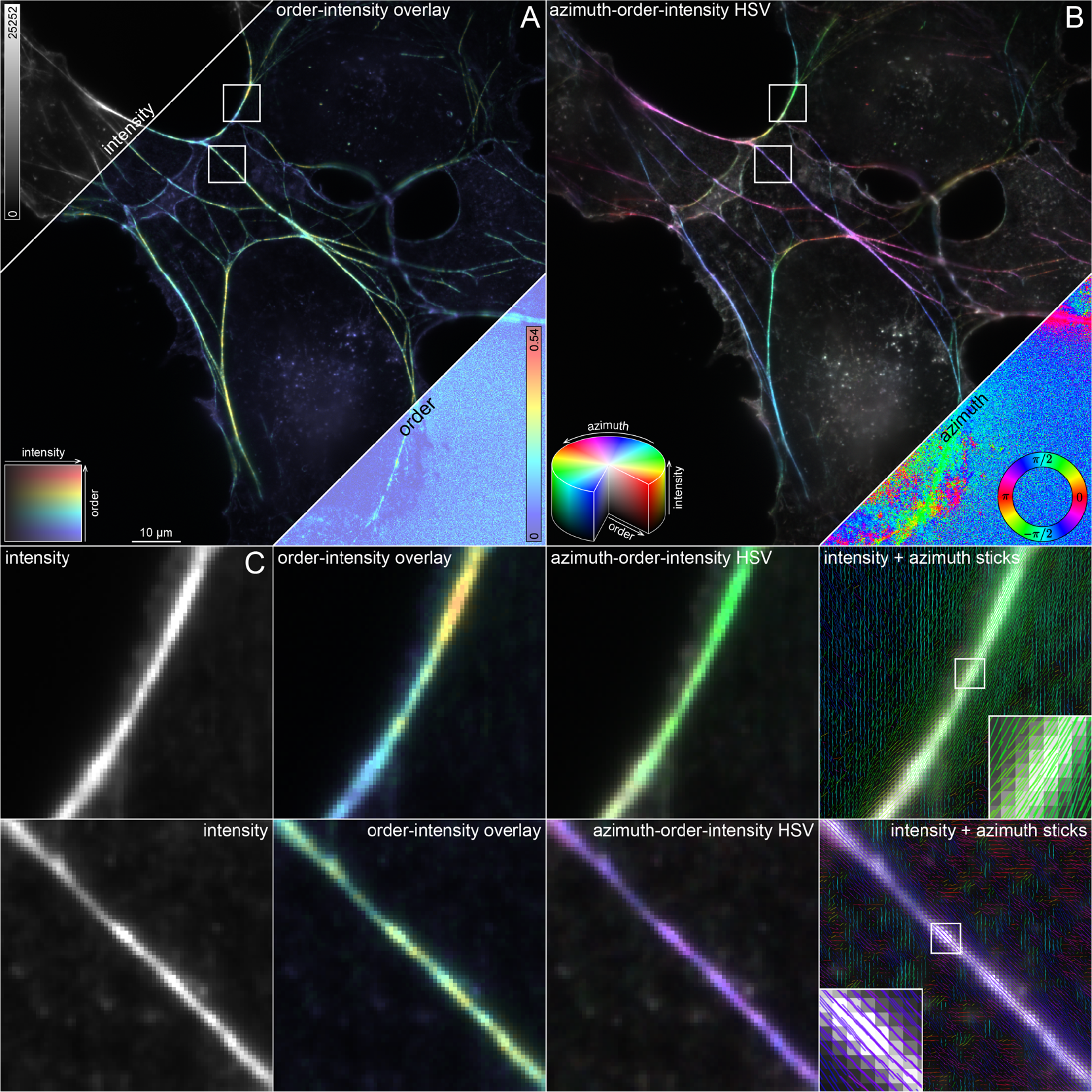
Image types and visualization options. Filamentous actin (F-actin) in COS-7 cells labelled with AF488-phalloidin and imaged with FPM to illustrate different output image types. (*A*) Order-intensity overlay (*middle*), made by combining the order (*lower right*) and intensity (*upper left*) images, with the latter acting as an opacity mask. (*B*) Azimuth-order-intensity HSV (*middle*), made by combining the azimuth (*lower right*), order (*A*), and intensity (*A*) images, which are used to set the hue, saturation, and value, respectively. (*C*) Magnified images of individual filaments indicated by the square ROIs in (*A*) and (*B*) showing—from left to right—the intensity, order-intensity overlay, azimuth-order-intensity HSV, and azimuth stick overlay. A small segment of each filament is highlighted with a square ROI and shown as a magnified inset to illustrate the expected alignment of the azimuths with the long axis of the filament.

The final two image types combine all three measurements—intensity, order, and azimuth—into a single visualization. For the azimuth-order-intensity HSV, data are converted into an HSV image with hue, saturation, and value (brightness) determined by the azimuth, order, and intensity, respectively (Fig 2B). In this way, both the order and orientation information are represented in a single image, with the brighter, more ordered regions appearing more prominent. Alternatively, users can display azimuths as sticks overlaid upon the intensity image, where the orientation and length of each stick correspond to the azimuth and order of its corresponding pixel (Fig 2C). The length, width, color, density, and transparency of the sticks can all be customized by the user. In this case, the colors of each stick correspond to the azimuth in each pixel. Both representations show that individual pixel azimuths are preferentially oriented parallel to the long axes of the filaments, as expected.

### Object-based analysis of punctate structures

To demonstrate key features of the software and the utility of an object-oriented analysis approach, we began by reanalyzing a previously published desmosome dataset (7). Desmosomes are intercellular adhesive junctions responsible for maintaining mechanical integrity in epithelia and cardiac muscle (19). We designed a series of EGFP-tagged constructs to study the desmosomal cadherin, desmocollin 2 (Dsc2a and Dsc2b), one of the membrane-spanning proteins that confer intercellular adhesion. The dataset comprises images of three unique Dsc2-EGFP chimeras which reveal the order and orientation of the Dsc2b ectodomain (ECTOb), Dsc2a ectodomain (ECTOa), and Dsc2a cytoplasmic domain (CYTO), along with a fourth construct where EGFP has been attached to the Dsc2a C-terminus with a flexible linker to act as a disordered control (LINK). There are two ECTOa groups, one of similar quality to the other groups—ECTOa (good)—and one of poor quality—ECTOa (poor)—which were collected in separate imaging sessions. ECTOa (poor), which suffers from low signal-to-background (S/B), would typically be excluded from biological analysis but is retained here to illustrate key software features.

All of the data in each group were processed and analyzed using default settings. Importantly, desmosomes are close to the diffraction limit (250 nm) in size and appear as either small puncta or slightly extended curvilinear structures in fluorescence microscopy. Therefore, the images were segmented using the built-in “Puncta” scheme, which employs traditional morphological operations followed by threshold selection with Otsu’s method (20) (S1 Text). Individual connected components in the mask are used to construct objects, each representing an individual desmosome and associated with an extensive set of extracted features which include FPM statistics derived from the polarization response data, morphological properties measured from the binary mask, and intensity information from the raw input data. Object features can be easily visualized and compared across and within groups, allowing for a more in-depth analysis than could be achieved using the FPM statistics alone. Moreover, the software allows users to label objects using property-based filtering or k-means clustering. This is particularly useful when objects display distinct morphological or intensity features and can be used to improve the quality of the underlying data prior to statistical comparisons.

One of the confounding factors that hinders precise and accurate determination of order and orientation parameters in FPM is the effect of noise on measurements of relative intensity changes (18). A distinctive feature of our software is the ability to estimate the effect of this noise locally by calculating the local S/B ratio for each object. In a typical image, the local S/B can vary considerably (21), even between objects that are relatively close to one another (Fig 3A, *left*). In low S/B regions, polarization-dependent intensity changes are dominated by noise, effectively lowering the measured order (Fig 3A, *right*). For instance, when examining two objects from the same cell border—one with low S/B (Fig 3B) and one with high S/B (Fig 3C)—the latter appears more ordered, despite representing the same construct and being localized to the same cell border. Likewise, the objects in ECTOa (good) appear considerably more ordered than those in ECTOa (poor) (Fig 3D), which suffer from low S/B (Fig 3E).

**Figure 3.**
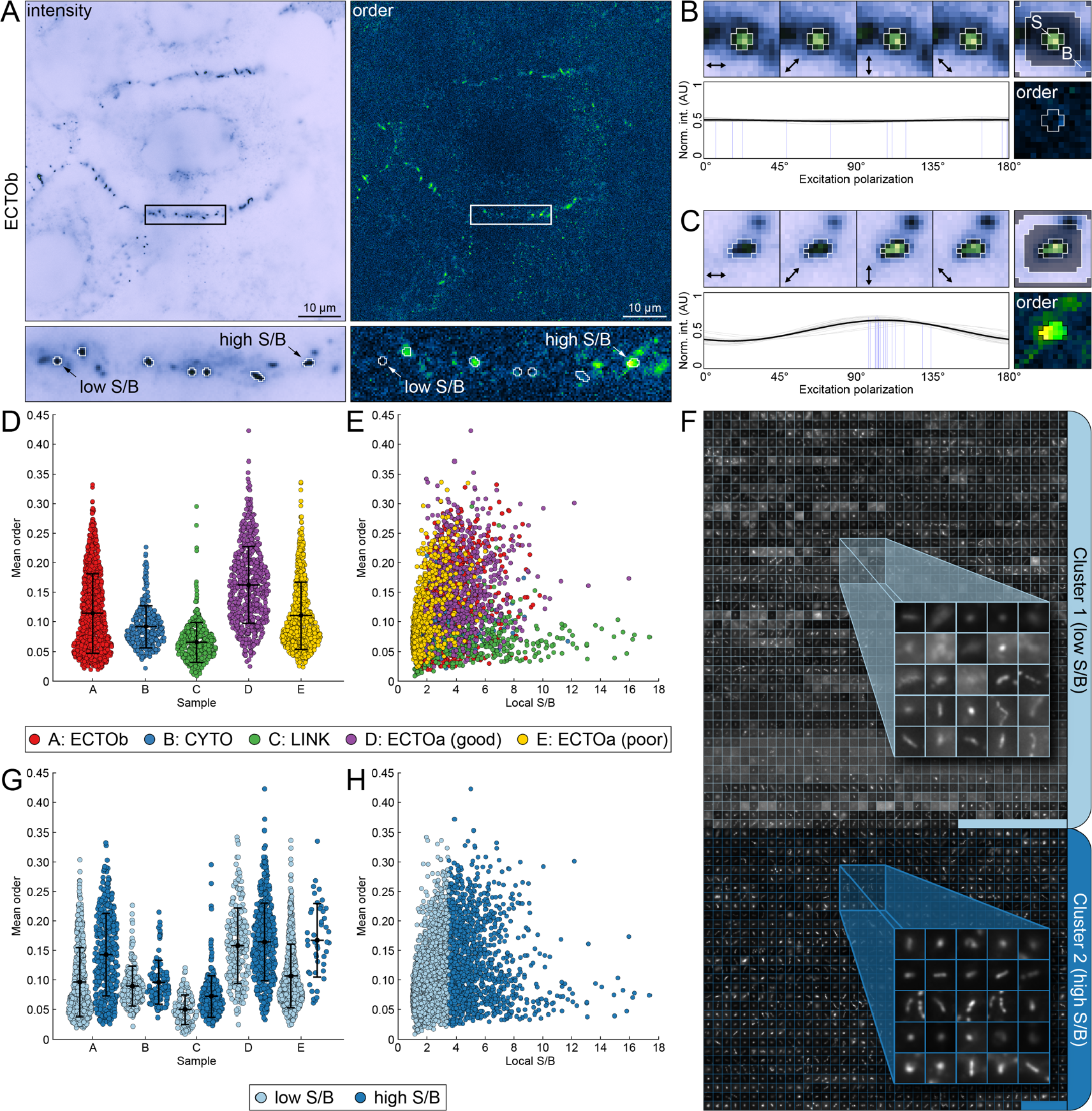
Object-based analysis of punctate structures. Desmosomal cadherin order probes expressed in A-431 cells, imaged with FPM, and analyzed with OOPS to demonstrate object-oriented FPM image analysis. (*A*) Representative intensity (*top left*) and order (*top right*) images of the Desmocollin 2b extracellular order probe (ECTOb). A single cell-cell border containing several desmosomes is indicated with a rectangular ROI and magnified below. Objects detected after segmentation are enclosed by white boundaries. Two objects that differ in order and signal-to-background ratio (S/B) are indicated by arrows. (*B*) Closer inspection of the low S/B object in (*A*). (*Top left*) Object intensity images at each excitation polarization, normalized to the maximum intensity in the stack. Arrows indicate the direction of the excitation field. (*Top right*) Average intensity image, normalized to the maximum intensity. Labels indicate regions used to determine local S/B (*S*: signal; *B*: background). (*Bottom left*) Pixel intensity stacks normalized to the total intensity and fit to a generic sinusoid (*gray*: individual pixel fits; *blue*: individual pixel azimuths; *black*: average of all fits). (*Bottom right*) Object order image. Pixels used to calculate mean order are enclosed in a white boundary. (*C*) Same as (*B*), but for the high S/B object indicated in (*A*). (*D*) Swarm plots showing mean order for each object in the desmosome dataset, grouped by construct: ECTOb (*red*), CYTO (*blue*), LINK (*green*), ECTOa (good) (*purple*), ECTOa (poor) (*yellow*). (*E*) Scatter plot of mean order versus local S/B for each object, grouped as in (*D*). (*F*) All objects across all groups were sorted into low and high S/B clusters using k-means clustering. Objects are represented by their average intensity images, which are tiled and stitched together within each cluster: Cluster 1 (*light blue*; low S/B) and Cluster 2 (*dark blue*; high S/B). (*G*) Same as in (*D*) but grouped by both construct and cluster. (*H*) Same as in (*E*) but grouped by cluster.

To examine in-depth the effect of S/B on measured order, we used k-means clustering to label the objects based solely on their local S/B, yielding two distinct clusters, one with low S/B (Cluster 1) and one with high S/B (Cluster 2) (Fig 3F) (S1 Text). Replotting the data with objects grouped by cluster (Fig 3G and Fig 3H) makes evident the effect of local S/B on observed order; in all cases, the mean order of the high S/B objects in Cluster 2 is greater than that seen in Cluster 1. Notably, when examining only objects in Cluster 2, the mean order of ECTOa (poor) is now remarkably close to that in ECTOa (good), despite differing considerably prior to clustering. Together, these data demonstrate the utility of our object-oriented approach; given the large number of object features extracted by the software, property-based labeling offers significant potential for enhancing workflows, revealing hidden relationships in the underlying data, and filtering out low quality data to enable more meaningful comparisons.

### Relative azimuth calculations reveal underlying molecular geometry

While order measurements yield important insights into how the labeled proteins are organized, they say little about the underlying geometry, which is better represented by the molecular azimuths. However, meaningful interpretation of these azimuths requires measuring from a suitable frame of reference—typically that of the object under investigation—which is unlikely to be aligned with the image itself. For eccentric, linear structures like F-actin filaments, this can be achieved by estimating the orientation of the azimuth sticks relative to the long axis of the object (Fig 2C). However, this is not quantitative and for smaller, non-linear objects, this procedure is cumbersome and time consuming. To illustrate this challenge, we consider a representative desmosome with an “S-shaped” morphology from the ECTOb dataset (Fig 4A, *top*). For each pixel in the object, the azimuth is simply the phase of a sinusoid fit to its polarization-dependent intensity profile (Fig 4A, *bottom*). In this context, the azimuth is an angle measured with respect to the direction of the excitation field in 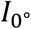 and therefore represents the average in-plane direction of the fluorescent dipoles relative to the horizontal direction in the image (*α*_image_). When the azimuths are plotted as sticks overlaid upon the average intensity image, they appear to be oriented perpendicular to the junction, yet this behavior is not reflected by the mean azimuth 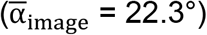 (Fig 4B) (S1 Text).

**Figure 4.**
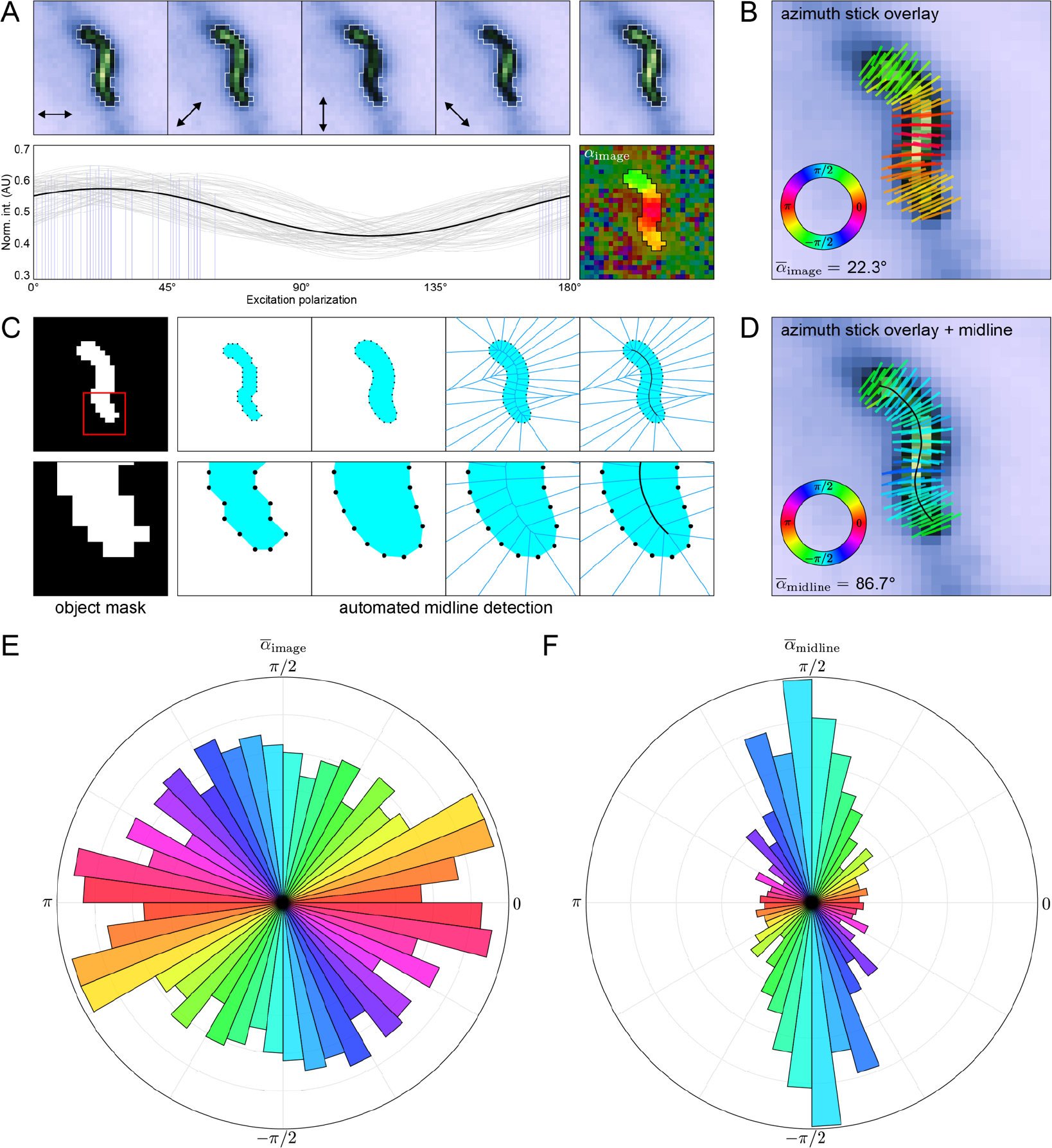
Relative azimuth calculations reveal underlying molecular geometry. (*A*) Representative “S-shaped” object from the ECTOb dataset. (*Top left*) Object intensity images at each excitation polarization, normalized to the maximum intensity in the stack. Arrows indicate the direction of the excitation field. (*Top right*) Average intensity image, normalized to the maximum intensity. (*Bottom left*) Pixel intensity stacks normalized to the total intensity and fit to a generic sinusoid (*gray*: individual pixel fits; *blue*: individual pixel azimuths; *black*: average of all fits). (*Bottom right*) Object azimuth image. Pixel values represent the angle of the azimuths with respect to the excitation field in 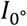 (*α*_image_). Background pixels are partially masked to highlight the object. (*B*) Average intensity image of the object in (*A*) with overlaid azimuth sticks, colored according to the direction, *α*_image_. (*C*) Simplified overview of the midline detection algorithm showing—from left to right—the binary mask defining the object; coordinates of the 8-connected perimeter pixels of the mask; boundary coordinates after dilation, smoothing, and linear arc interpolation; Voronoi diagram of the adjusted boundary points; and final midline detected from the central most edges of the Voronoi diagram after smoothing and interpolation. (*D*) Same as in (*B*), but with the midline overlaid and azimuth sticks colored according to their direction relative to the nearest midline tangent (*α*_midline_). (*E*) Polar histogram showing the distribution of 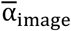 for all objects in the ECTOb dataset. Object 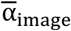 values initially in the range [−π/2, π/2] are duplicated and shifted by π to show each pair of equivalent, opposite directions. (F) Same as in (E), but for object 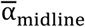 directions.

To facilitate quantification of azimuth orientations with respect to the objects themselves, we developed an automated midline detection algorithm (Fig 4C) (S1 Text). In brief, we start with a set of points representing the 8-connected boundary of the object mask which is dilated, smoothed, and then used to construct a Voronoi diagram, the central most edges of which comprise the object midline. The midline points are extracted and smoothed, and the local orientation of the object is estimated by assigning to each pixel the value of the nearest midline tangent. Finally, azimuths are recalculated with respect to the orientation of the midline (α_midline_). For the S-shaped desmosome, the mean azimuth measured with respect to the midline more accurately reflects the qualitative appearance 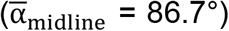 (Fig 4D). To demonstrate the utility of this approach, we compared azimuth calculation methods for all objects in the ECTOb dataset. When not adjusted to account for the orientation of the objects, azimuths appear randomly distributed and provide little information about the underlying geometry (Fig 4E). On the other hand, when measured with respect to the midlines, azimuths are overwhelmingly perpendicular to the objects (Fig 4F). This reveals a consistent organization of the labeled molecules with respect to their larger complexes, while also suggesting the individual fluorescent dipoles are arranged symmetrically about the membrane normal. Importantly, this information would be entirely lost if azimuths were not adjusted to account for the orientation of the objects, highlighting the power and utility of our approach.

### Object-based analysis of filamentous structures

The object-oriented analysis approach is fully compatible with any sample morphology, given an appropriate mask. Here, to demonstrate the flexibility of the approach, we used OOPS to analyze the orientation of the F-actin cytoskeleton in HUVECs grown either in static conditions or under fluidic shear stress (FSS). In cells grown under static conditions, the filaments appear randomly oriented (Fig 5A), whereas those in the cells grown under FSS appear to be preferentially oriented along the direction of the flow (Fig 5B). For both sets of samples, images were segmented using the built-in “Filaments” scheme, designed to detect linear, extended structures (S1 Text). In both groups, the azimuths are oriented along the direction of the filaments and therefore report the directions of the filaments themselves (Fig 5C and 5D). For cells grown statically, azimuths are randomly oriented with respect to the image, consistent with the appearance of the filaments. On the other hand, for cells grown under FSS, the azimuths are roughly aligned with the horizontal direction in the image, consistent with a reorganization of the actin cytoskeleton.

**Figure 5.**
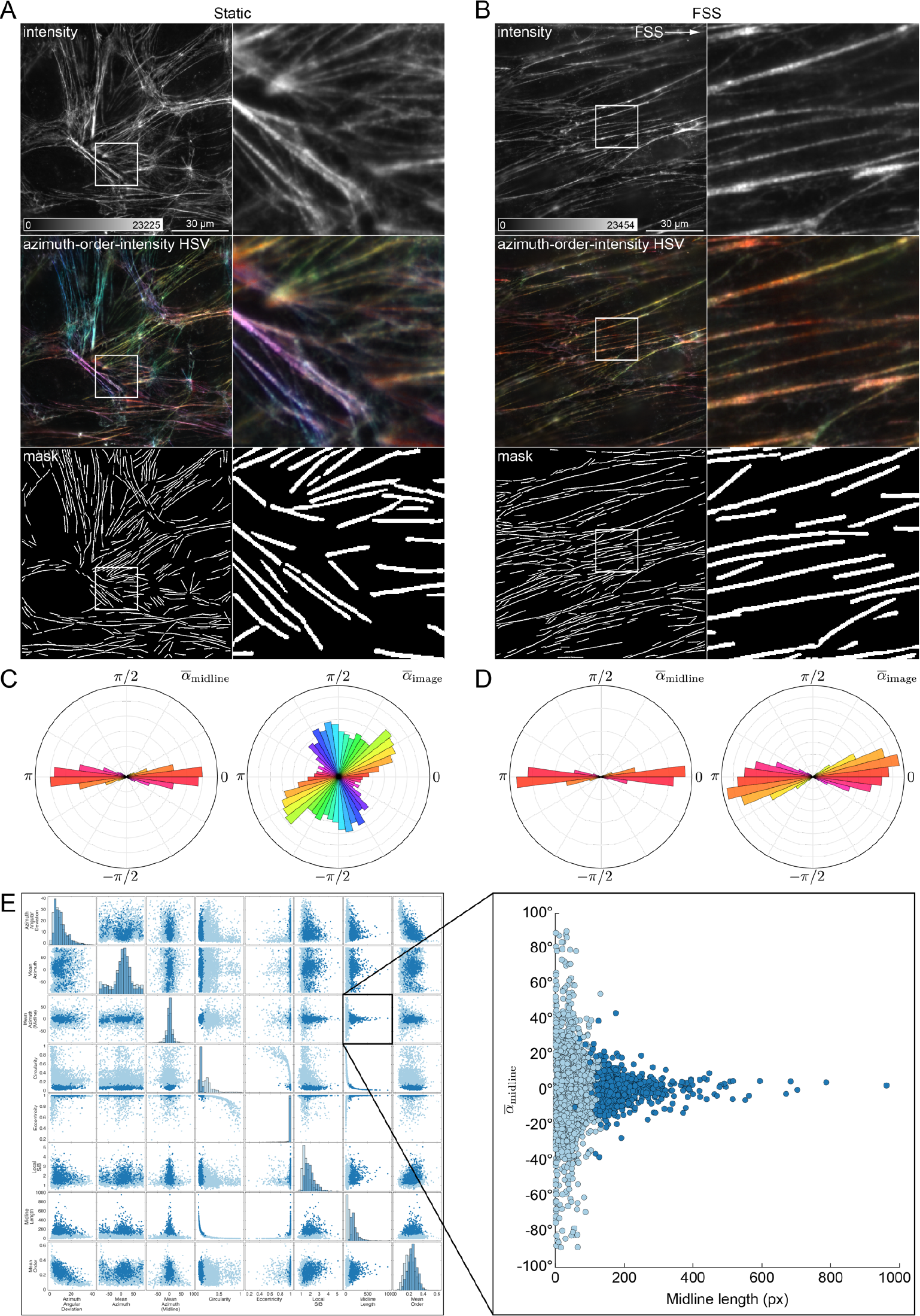
Object-based analysis of filamentous structures. (*A*) Human umbilical vein endothelial cells (HUVECs) grown in static conditions, labelled with AF488-phalloidin, and imaged with FPM. From top to bottom: average intensity image, azimuth-order-intensity HSV image, and binary mask showing locations of detected filaments. The region indicated by a square ROI is shown magnified to the right. (*B*) Same as (*A*) but for HUVECs grown under fluidic shear stress (FSS). White arrow indicates the flow direction. (*C*) Polar histograms showing 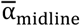 (*left*) and 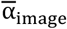 (*right*) distributions for filaments in cells grown statically. (*D*) Same as (*C*), but for cells grown under FSS. (*E*) Filaments were clustered into two groups based on their morphology and intensity features. (*Left*) Scatter plot matrix showing relationship between several morphology, intensity, and FPM statistics for all filaments, colored by cluster label. (*Right*) Magnified scatter plot of 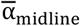 versus midline length, showing the difference in 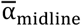 between long and short filaments.

To extract additional information, we labeled the segmented filaments using k-means clustering, informed this time by a large set of morphological and intensity features (S1 Text) (Fig 5E). Interestingly, the filaments fell into two clusters, differing mostly in terms of their size and relative intensity. The shorter, dimmer filaments were characterized by average azimuths that were not well-aligned with the object midlines, while the longer, brighter filaments displayed azimuths typical of F-actin labeled with AF488-phalloidin. Importantly, the FPM-derived order and orientation statistics were not used to inform the clustering, suggesting the differences observed therein are related to the morphology of the filaments. In that sense, clustering or property-based labeling presents a straightforward means to reveal subtle, otherwise hidden relationships in the underlying data. This procedure could also be used to refine the collection of objects by removing those deemed undesirable for analysis.

## Conclusions

This study introduces OOPS as a flexible, GUI-driven MATLAB package for the analysis of molecular order and orientation in FPM images. It provides an intuitive platform for object-based image analysis, enabling users to uncover meaningful, otherwise hidden features in the underlying data. OOPS can be adapted to a variety of acquisition setups and is agnostic to both sample geometry and labeling method, as demonstrated here via the analysis of punctate and filamentous structures.

## Availability and future directions

OOPS is freely available under the GNU GPL 3.0 license and can be downloaded at https://github.com/Mattheyses-Lab/OOPS. All of the data analyzed in this study are included with the software. Descriptions of each dataset, as well as instructions to reproduce the analyses shown are given in S1 Text. A detailed user manual is included with the software and in S2 Text. We will continue to implement new functionality as our own analysis requirements evolve, and we encourage users to request features that would enhance the capabilities of the software.

## Author contributions

Desmosome sample preparation and data collection: WFD

F-actin sample preparation: TJN

F-actin data collection: RMA and WFD

Data analysis: WFD

Software design and programming: WFD

Software testing: WFD, TJN, and RMA

Manuscript and figures: WFD and ALM

Funding acquisition: ALM

